# Extracellular vesicles carrying HIV-1 Nef induce trained immunity in myeloid cells

**DOI:** 10.1101/2022.06.24.497538

**Authors:** Larisa Dubrovsky, Beda Brichacek, N.M. Prashant, Tatiana Pushkarsky, Nigora Mukhamedova, Dragana Dragoljevic, Michael Fitzgerald, Anelia Horvath, Andrew J. Murphy, Dmitri Sviridov, Michael I Bukrinsky

## Abstract

**Summary:** Persistent inflammation is a hallmark of HIV infection and is not reversed after suppression of viral replication by anti-retroviral therapy (ART). One explanation for chronic inflammation in ART-treated HIV-infected individuals is hyperreactivity of the myeloid cells due to a phenomenon called ‘trained immunity’. Here, we demonstrate that human monocyte derived macrophages originating from monocytes initially treated with extracellular vesicles containing HIV-1 protein Nef (exNef), but differentiating in the absence of exNef, released increased levels of pro-inflammatory cytokines after lipopolysaccharide stimulation. This effect was associated with epigenetic changes related to inflammation and cholesterol metabolism pathways, upregulation of the lipid rafts, and was blocked by methyl-β-cyclodextrin, statin, and inhibitor of activity of lipid raft-associated receptor IGF1R. Bone marrow-derived macrophages from exNef-injected mice had higher abundance of lipid rafts and produced elevated levels of TNFα. These phenomena are consistent with trained immunity and may contribute to persistent inflammation and co-morbidities in HIV-infected individuals.

## Introduction

Combination anti-retroviral therapy (cART) has dramatically altered the course and prognosis of HIV infection, changing it from a fatal condition to a manageable chronic disease (Tseng et al., 2015). However, despite a significantly increased life expectancy, people living with HIV (PLWH), even those who consistently maintain undetectable viral load, remain at increased risk of developing a range of co-morbidities that affect both longevity and the quality of life of this population presenting a major clinical problem (Lerner et al., 2019). These co-morbidities, which include cardiovascular disease (CVD), type II diabetes and neurocognitive dysfunction (HAND), have one common feature underlying their pathogenesis – persistent low-grade inflammation (Hileman and Funderburg, 2017). Indeed, PLWH are characterized by persistent increase in inflammatory markers and chronic immune activation, which are mitigated by ART but do not reverse to levels measured in HIV-uninfected population (Lederman et al., 2013; Zicari et al., 2019). The reason for this phenomenon has been primarily attributed to bacterial leakage due to disruption of the tight junctions in the intestinal epithelium, which may not be fully restored by ART (Nazli et al., 2010). Another potential cause for persistent inflammation may be the continuous release of HIV-related pro-inflammatory factors, in particular Nef protein, from the viral reservoirs (Hileman and Funderburg, 2017; Raymond et al., 2019; Stevenson et al., 2021). Nef is released from HIV-infected cells predominantly as a component of exosomes (exNef) (Campbell et al., 2008; Ellwanger et al., 2017; Lenassi et al., 2010; McNamara et al., 2018; Puzar Dominkus et al., 2017), and exNef was shown to exert a potent pro-inflammatory effect by suppressing cholesterol efflux and elevating the lipid raft abundance on myeloid cells (Mukhamedova et al., 2019).

Yet another possible contributor to the observed long-term immune activation occurring on the background of successful ART might be a “legacy effect”, when an exposure to a pathogen produces effects that last long after the pathogen is eliminated. One mechanism of this phenomenon is ‘trained immunity’ (Crisan et al., 2016; Netea et al., 2011; Quintin et al., 2014). This paradigm, introduced by Netea and colleagues a decade ago (Netea *et al*., 2011), provides an elegant explanation for the long known phenomenon of immunologic memory associated with the innate immune responses, exemplified by ‘non-specific’ protection provided by live vaccines against infections unrelated to the vaccine immunogen (de Bree et al., 2018; Garly et al., 2003; Weinstein et al., 2010). The mechanism of this effect was shown to be metabolic-epigenetic, i.e., involve changes in cell metabolism leading to a modification of the chromatin composition in myeloid cells occurring in response to an infectious agent and resulting in increased expression of pro-inflammatory genes after a subsequent stimulus with an unrelated pathogen (van der Heijden et al., 2018). The factors shown to induce trained immunity include microbial and non-microbial stimuli, such as LPS, β-glucan, oxLDL, lipoprotein(a), aldosterone (Bekkering et al., 2014; Geng et al., 2016; Neidhart et al., 2019; Riksen and Netea, 2020; van der Heijden et al., 2020; van der Valk et al., 2016). The present study was undertaken to investigate whether exposure to Nef, the key pathogenic factor of HIV (Deacon et al., 1995; Hanna et al., 1998; Kestler et al., 1991; Kirchhoff et al., 1995), leaves a memory in monocyte-derived macrophages that may contribute to increased inflammatory responses to subsequent stimulation.

## Results

### Trained immunity induced by exNef

To investigate the ability of Nef to induce trained immunity, we used a classical approach introduced by the Netea’s group (Bekkering et al., 2016). Isolated human primary monocytes were exposed for 48 h to an agent (in our case, Nef), washed and left in the culture medium for 6 days to differentiate into monocyte-derived macrophages (MDM), which then are restimulated for 4 h with an unrelated stimulus, LPS (Fig. 1A). The change in cytokine production after stimulation, relative to cells not exposed to the training agent, is taken as a marker of trained immunity. Given that Nef is released from HIV-infected cells predominantly within the extracellular vesicles (EVs) (Campbell *et al*., 2008; Ellwanger *et al*., 2017; Lenassi *et al*., 2010; McNamara *et al*., 2018; Puzar Dominkus *et al*., 2017), we used EVs produced by the Nef-transfected HEK293T cells (exNef) as a source of Nef; EVs from RFP-transfected cells (exCont) served as controls. These EVs were characterized in our previous studies and shown to exhibit the size and markers characteristic for exosomes (Dubrovsky et al., 2020). Results presented in Fig. 1B demonstrate increased production of TNFα and IL-6 by MDM that differentiated from monocytes exposed to exNef relative to those exposed to exCont.

**Figure 1.**
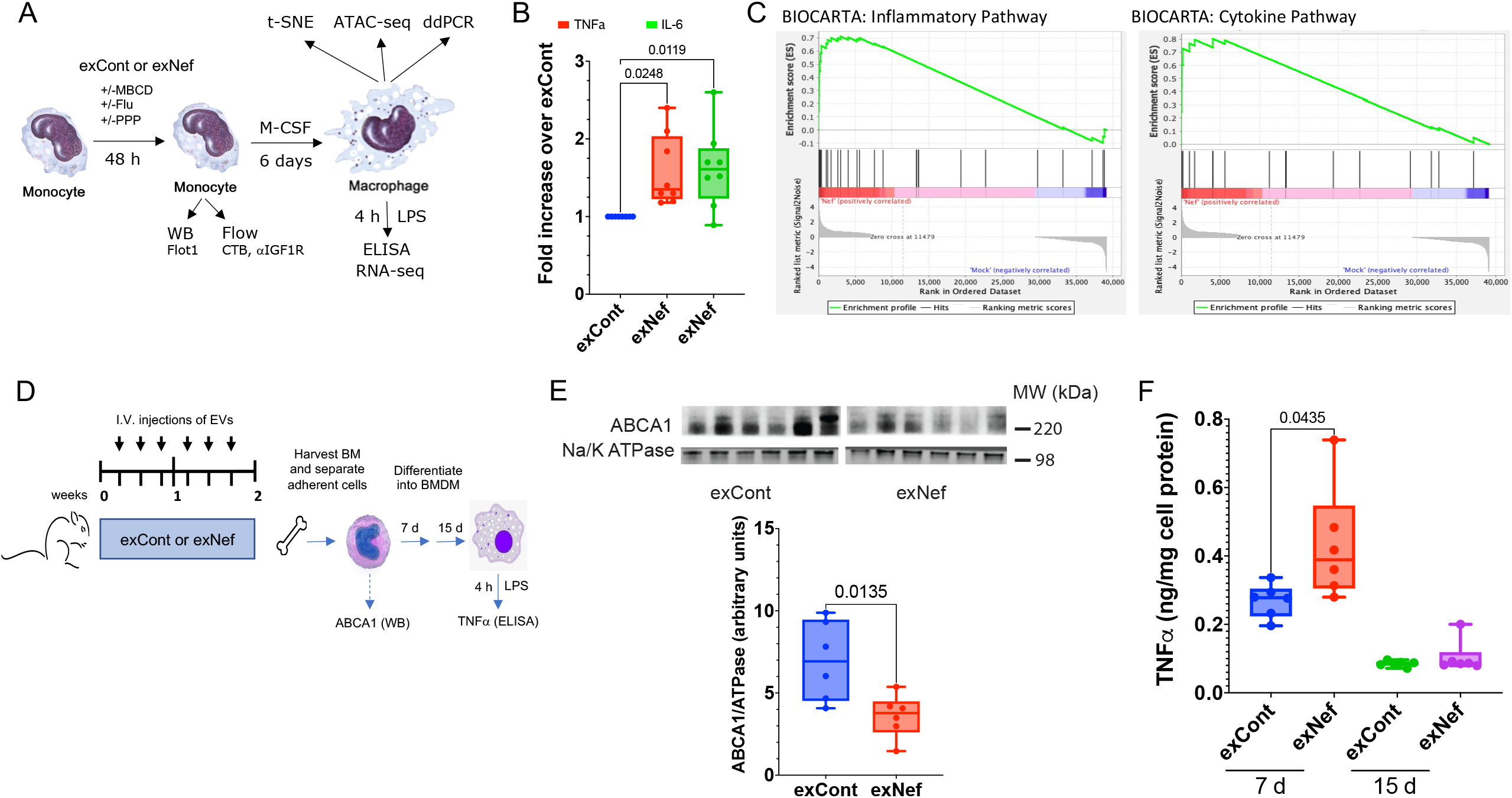
Analysis of trained immunity. **A** – Diagram of *in vitro* experiments performed in this study. Monocytes treated for 48 h with exNef or exCont in the presence or absence of methyl-β - cyclodextrin (MBCD), Fluvastatin (Flu), or picropodophyllin (PPP) were analyzed by Western blotting (WB) for flotillin 1 (Flot1) or by flow cytometry for lipid rafts (staining with CTB) and IGF1R. After washing, cells were incubated for 6 days in the presence of M-CSF and characterized by t-SNE for activation markers or by ddPCR for gene expression. Following a LPS stimulation, gene expression was characterized by RNA-seq, and cytokine production measured by ELISA. Comparisons were made between cells treated with exNef and exCont. **B** – Analysis of TNFα and IL-6 production by LPS-treated macrophages exposed to exNef or exCont. The graph shows fold increase of cytokine production by exNef-treated over exCont-treated cells from 8 donors, analyzed by Friedman multiple comparison test with Dunn’s correction (Prism 9.0). P values are shown above bars. **C** – Gene Set Enrichment Analysis (GSEA) (Subramanian *et al*., 2005) showing normalized enrichment scores for genes differentially expressed between exNef and exCont and participating in the inflammatory response (top) and cytokine pathway (bottom); the gene-sets were obtained from BioCarta (Nishimura, 2001). **D** – Diagram of *in vivo* experiments performed in this study. Mice were injected with exCont or exNef for 2 weeks, adherent cells from the bone marrow were isolated, analyzed for ABCA1, and cultured for 7 or 15 days without EVs. Following LPS stimulation, TNFα was measured by ELISA. **E** – Upper panel – Western blot of ABCA1 and Na/K ATPase (loading control) of cell lysate of BMDM from individual mice; bottom panel – densitometric analysis of the gel above (abundance of ABCA1 relative to Na/K ATPase, arbitrary units). P value was calculated by unpaired two-tailed parametric test. **F** – TNFα produced by BMDM stimulated with LPS on days 7 or 15. P value (only significant one is shown) was calculated by unpaired two-tailed parametric test.

To assess the genome-wide gene-expression changes associated with exNef treatment, we performed whole transcriptome RNA sequencing (RNA-seq) and gene expression analysis of exNef-trained MDM from two donors. Gene Set Enrichment Analysis (GSEA (Subramanian et al., 2005)) showed that the most upregulated gene sets were enriched in genes involved in cytokine and inflammatory pathways (Fig. 1C). These results were reproduced with MDM from 8 more donors (Fig. S1). Again, exNef treatment of monocytes altered transcriptional response to LPS of differentiated MDM (Fig. S1A), with gene set enrichment analysis identifying genes in the cytokine and inflammation pathways as the most affected by the treatment (Fig. S1B). Furthermore, leading edge analysis of the genes deregulated across the samples from 8 different donors showed that a number of cytokine genes, including *IL-2, IL-3, IL-6, IL-11, IL-10, IL-18, CXCL8*, and *TNFα*, drive the activation of several inflammation-related pathways (Fig. S1C).

To determine whether trained immunity observed with monocytes treated with exNef *in vitro* also occurs *in vivo*, we intravenously injected either exNef or exCont (EVs produced by *GFP*-transfected HEK293T cells) into C57BL/6J mice for 2 weeks, isolated bone marrow progenitor cells, differentiated them into BMDM, and tested responses to LPS stimulation (Fig. 1D). Analysis of isolated bone marrow cells demonstrated lower abundance of ABCA1 in cells from exNef treated animals (Fig. 1E), an expected effect of exNef. Isolated cells were plated on dishes and allowed to differentiate into BMDM in the absence of EVs for 7 and 15 days. On day 7 of differentiation, plasma membranes isolated from BMDM of animals treated with exNef had significantly higher abundance of lipid rafts (determined as incorporation of exogenous [^3^H]cholesterol and abundance of flotillin-1 in gradient fractions corresponding to the lipid rafts, fractions 2-4, Fig. S2) than membranes from cells of animals treated with exCont. Correspondingly, in response to LPS stimulation, BMDM from mice treated with exNef produced significantly more TNFα than cells from exGFP-injected mice at a 7-day time point (Fig. 1F). Cells stimulated with LPS on day 15 produced very little TNFα and differences between cells from exCont and exNef treated animals were not significant. This finding suggests that exNef modifies bone marrow myeloid progenitor cells to produce hyperresponsive macrophages.

### Characterization of trained MDM

Following differentiation into MDM, cells were analyzed by multi-color flow cytometry. The cell markers in this analysis represented those commonly used to characterize the phenotype of monocyte/macrophages: CD14, CD16, CD-11b, CD40, CD163, CD38, CD80, CD64, HLA-DR, CD68. These markers are associated with the role of macrophages in pro- and anti-inflammatory responses, although the precise assignment of a certain marker to a certain phenotype remains controversial (Atri et al., 2018; Kapellos et al., 2019). Treatment with exNef resulted in significant changes in the distribution of the cells between the populations characterized by levels of marker expression presented as t-distributed stochastic neighbor embedding (t-SNE) plot (Fig. 2A). All cell populations enriched by exNef treatment (over 50% increase in cell numbers) were characterized by low expression of CD16 (peaks shifted to the left of the diagram), and most had high expression of CD80, CD64, CD68 and HLA-DR (Fig. 2B). The CD16^low^ phenotype is characteristic for M1 macrophages (Gui et al., 2012). Consistently, pro-inflammatory markers on a number of cell populations were upregulated. Analysis of cells from 2 additional donors confirmed the effect of exNef on expression of inflammation-associated markers (Fig. S3A,C). Again, CD16 expression was low on cells from most populations enriched after exNef treatment. Although specific patterns of the changes somewhat differed between the donors, high expression of CD40 and CD64, and CD11b was noted in several populations of exNef-treated cells from donors B and C (Fig. S3B,D). These results indicate that treatment with exNef altered the differentiation of monocytes, promoting the pro-inflammatory phenotype.

**Figure 2.**
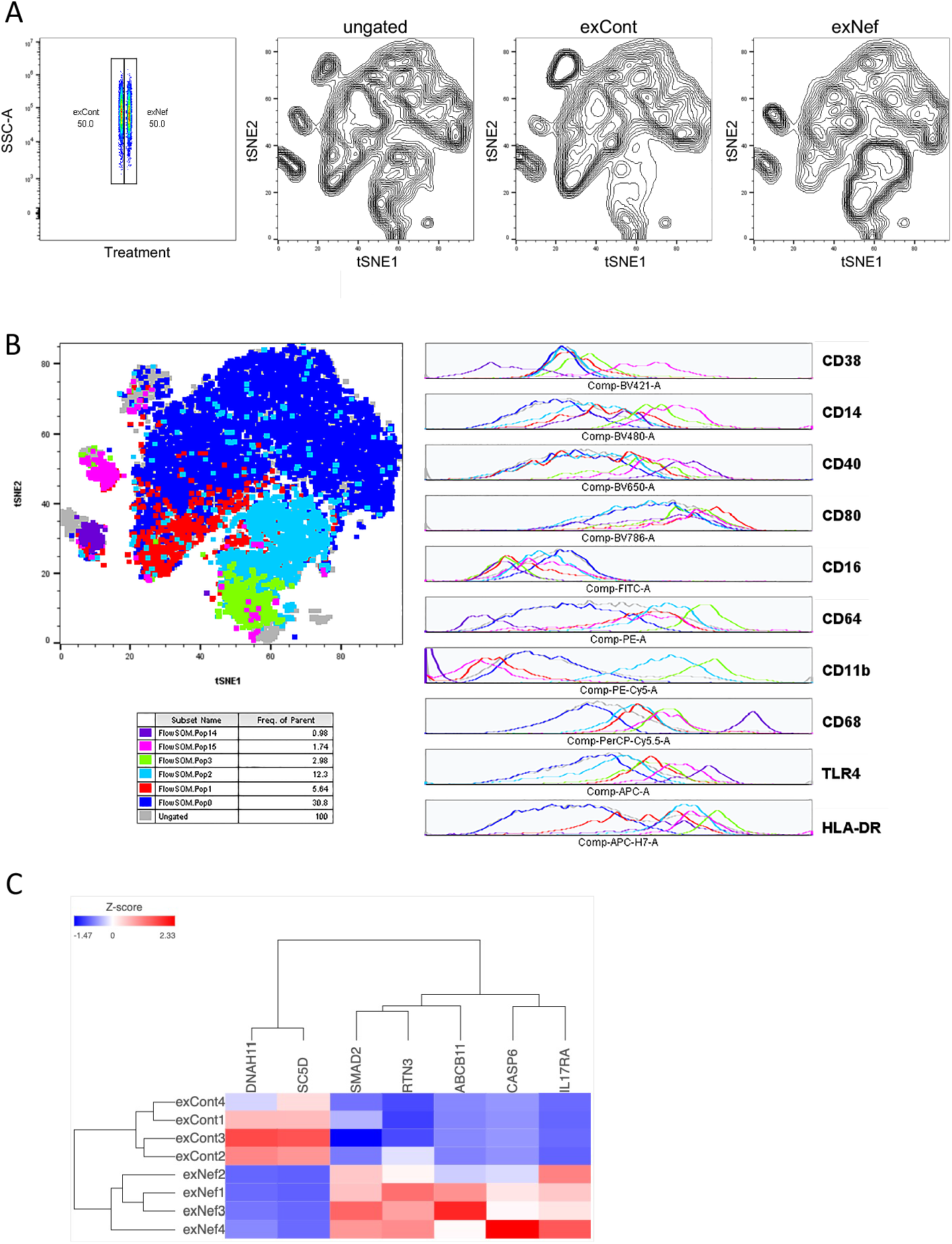
Characterization of trained MDM. **A** - Equal amounts of one donor’s MDM exposed to exCont or exNef were subjected to t-SNE (t-stochastic neighbor embedding) analysis. Unbiased number of clusters (populations) were determined by Phenograph technique and applied to FlowSom analysis to visualize cytometry data. Results are shown for ungated population, and after gating on exCont or exNef exposed cells. **B** - Phenotypic outcome was evaluated in the populations where exNef-treated cells prevailed over exCont-treated by 50% or more. Top left panel shows distribution of these cell populations, and the table underneath presents the frequency of parent population for each cluster. The diagrams on the right show expression of indicated markers on the cells of each color-coded population. **C** - The heatmap shows genes which localization changed between open and closed chromatin regions as a result of exNef treatment. Results are shown for cells from 4 donors treated with exCont or exNef.

Previous studies demonstrated the key role of epigenetic changes in the trained immunity phenotype (Fanucchi et al., 2021). To determine whether exNef induced epigenetic modifications in monocytes, we employed Assay for Transposase-Accessible Chromatin using sequencing (ATAC-seq) to reveal changes in genes’ localization to the open (active) chromatin regions. Results presented in Figure 2C for cells from 4 donors show significantly affected genes that changed localization between open and closed chromatin as a result of exNef treatment. These genes belonged to two pathways, cholesterol metabolism and inflammation (Table S1). Again, substantial inter-donor variability was observed in identity of the genes affected by exNef treatment, suggesting a significant effect on trained immunity of individual-specific characteristics (age, gender, inflammatory status, etc). These findings indicate that changes in monocyte differentiation and inflammatory response were accompanied by epigenetic modifications, consistent with mechanism of trained immunity.

### Training is sensitive to inhibition of cholesterol biosynthesis and IGF1R signaling

Previous study of the β-glucan-induced trained immunity demonstrated that the molecular mechanism behind this phenomenon involves increased production of mevalonate, an intermediate product of cholesterol biosynthesis, its secretion, and autocrine stimulation of the insulin-like growth factor-1 receptor (IGF1R), signaling from which greatly potentiated training (Bekkering et al., 2018). Treatment with statin during the training abolished the trained immunity (Bekkering *et al*., 2018), demonstrating the critical role of cholesterol biosynthesis pathway in trained immunity. To determine if similar mechanisms operate in the trained immunity induced by exNef, we analyzed expression of several cholesterol biosynthesis rate-limiting genes in MDM exposed to exNef or exCont. This analysis revealed upregulation of HMGCR (3-Hydroxy-3-Methylglutaryl-CoA Reductase) in cells treated with exNef (Fig. 3A). No significant changes were detected in expression of two other tested cholesterol biosynthesis genes, SQLE (squalene epoxidase) and MVK (mevalonate kinase) (Fig. 3A). Cholesterol biosynthesis (measured by assessing incorporation [^3^H]acetate into cholesterol) in monocytes is very low (Fernandez-Ruiz et al., 2016), preventing reliable detection of significant differences between exCont- and exNef-treated cells. However, when differentiated MDM were treated with exNef, they showed increased rate of cholesterol biosynthesis, relative to cells treated with exCont (Fig. 3B). This finding is consistent with previously demonstrated activation of cholesterol biosynthesis genes’ expression by endogenously expressed Nef (van ‘t Wout et al., 2005). The role of cholesterol biosynthesis induction in training was supported by the inhibitory effect of an inhibitor of HMGCR, Fluvastatin, on exNef-induced training. Pre-treatment of cells with Fluvastatin reduced TNFα production after LPS stimulation to the levels observed in exCont-treated cells (Fig. 3C).

**Figure 3.**
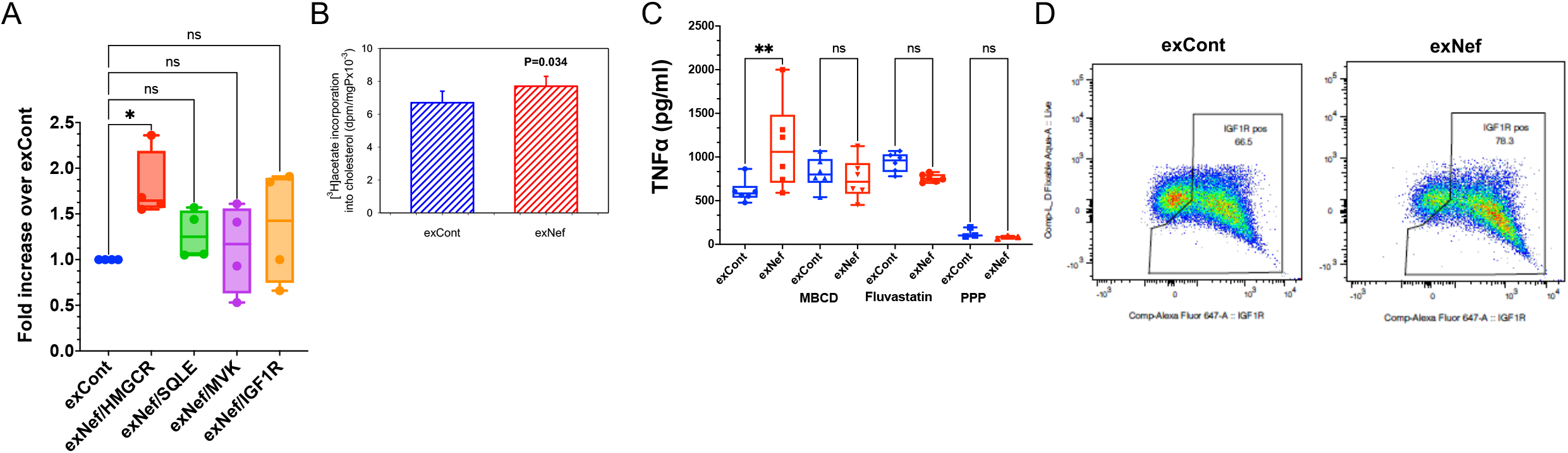
Mechanisms of exNef-induced training. **A** - Results of ddPCR analysis of expression of *MVK, SQLE, HMGCR* and *IGF1R* genes in monocytes treated with exCont or exNef (24 h after removal of EVs). Bars show fold increase of gene expression in exNef-treated relative to exCont-treated cells. Analysis was performed in cells from 4 donors, and results were analyzed by ANOVA with Dunnett correction for multiple comparisons. **B** - Incorporation of [^14^C]acetate into cholesterol after 2 h incubation with MDM treated with exCont or exNef. Bars show means ± SD of dpm per mg of total cell protein from quadruplicate determinations. P value was calculated by unpaired two-tailed parametric test. **C** – ExNef-induced upregulation of TNFα production by LPS-treated MDM was reversed by methyl-β-cyclodextrin (MBCD), Fluvastatin (Flu), and picropodophyllin (PPP). Flu and PPP were added together with EVs, MBCD was added after washout of EVs. Results were analyzed by one-way ANOVA, with Holm-Sidak’s correction for multiple comparisons. **p=0.0024. **D** – IGF1R presentation on MDM exposed to exNef and exCont. Monocytes were analyzed after washout of EVs.

Similar to observations with β-glucan-induced trained immunity (Bekkering *et al*., 2018), exNef-induced training was also blocked by picropodophyllin (PPP), an inhibitor of IGF1R signaling (Fig. 3C). However, in contrast to β-glucan, exNef increased the presentation of IGF1R on the membrane of treated cells (Fig. 3D). Interestingly, exNef did not increase the mRNA expression of IGF1R (Fig. 3A), suggesting that a post-transcriptional mechanism was responsible for increased IGF1R presentation on the membrane. Thus, training induced in monocytes by exNef had properties similar to immune training described by Netea and colleagues (Bekkering et al., 2021).

### The role of lipid rafts in exNef-induced trained immunity

ExNef has been shown to increase the abundance of lipid rafts on target cells, promoting activity of lipid raft-associated receptors (Mukhamedova *et al*., 2019). Given that IGF1R is associated with lipid rafts (Delle Bovi et al., 2019; Hong et al., 2004; Huo et al., 2003; Romanelli et al.,

2009; Sural-Fehr et al., 2019), exNef-mediated training may be controlled at the level of IGF1R presentation at the lipid rafts. Indeed, exNef increased the staining of the monocytes with CTB, which specifically binds to the lipid raft marker GM1 (Fig. 4A). This result was confirmed by analysis of the lipid raft component flotillin-1 in the plasma membrane of the cells treated with exNef and exCont: significantly higher abundance of flotillin-1 was detected in exNef-treated cells (Fig. 4B,C). The role of the lipid rafts in exNef-induced training was further supported by the reversal of the training-specific increase of TNFα production by treatment of the monocytes with methyl-β-cyclodextrin (MβCD) (Fig. 3C), which disrupts lipid rafts by extracting cholesterol.

**Figure 4.**
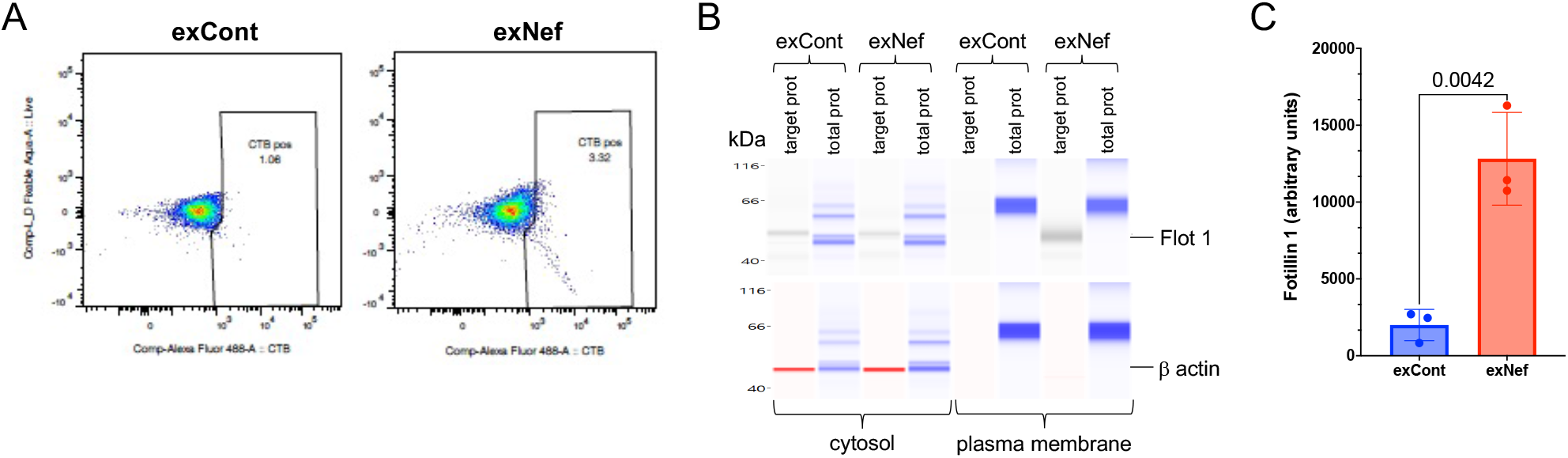
ExNef upregulates lipid rafts. **A** – CTB staining of lipid rafts in monocytes treated with exNef or exCont. Monocytes were analyzed after washout of EVs. **B** – Flotillin 1 was analyzed in plasma membranes isolated from monocytes after washout of EVs following treatment with exCont or exNef. Total protein staining provided loading control. **C** – Bars show mean±SD of gel area corresponding to Flotillin 1 band adjusted to total protein (Jess instrument software), calculated for cells from 3 donors. P value was calculated by unpaired two-tailed t-test.

Taken together, these findings suggest that exNef may induce trained immunity by stimulating cholesterol biosynthesis and increasing the abundance of the lipid rafts, thus increasing presentation of IGF1R and its signaling.

## Discussion

In this study, we describe how treatment of monocytes with extracellular vesicles carrying HIV-1 protein Nef (exNef) induces a pro-inflammatory memory in macrophages that differentiate from treated monocytes. Our findings demonstrate enhanced secretion of two key cytokines, TNFα and IL-6, increased expression of pro-inflammatory genes and changes in chromatin composition, suggesting epigenetic alterations induced by exNef. These effects of exNef on monocytes are similar to those of the fungi cell wall polysaccharide β-glucan, as described by Quintin et al (Quintin et al., 2012) and termed ‘trained immunity’ (Netea et al., 2020).

Phenotypic analysis of MDM produced from monocytes exposed to exNef revealed formation or enrichment of new cell populations characterized by expression of several markers associated with the M1 polarization. However, none of these populations expressed a classical M1 polarized phenotype, consistent with previous studies that demonstrated that training of monocytes does not induce classical M1 or M2 macrophage phenotypes (Bekkering *et al*., 2014; Quintin *et al*., 2012). To identify genes whose promoters were associated with open chromatin regions and thus were more sensitive to activation stimuli, we relied on ATAC-seq. Previous studies detected activation of multiple genes belonging to metabolic and inflammatory pathways in β-glucan-trained MDM (Bekkering *et al*., 2016; Netea *et al*., 2020; Novakovic et al., 2016; Riksen and Netea, 2020; van der Heijden *et al*., 2018). Consistent with these reports, our analysis of MDM trained by exNef revealed enrichment in the open chromatin regions of genes associated with inflammation, immune response, and insulin metabolism; activation of these pathways may contribute to the co-morbidities in PLWH. Consistent with functional manifestations of trained immunity, MDM trained with exNef responded to LPS stimulation with increased expression levels of a number of genes, including IL-3, IL-6, IL-9, IL-11, CXCL8, CSF3, CSF2, and TNFα, which participate in multiple pathways, including cytokine, inflammation and proteasome pathways. The mechanism behind exNef-induced trained immunity is also consistent with that described for β-glucan (Bekkering *et al*., 2018): it depends on activation of cholesterol biosynthesis and IGF1R signaling. Another important component of exNef-induced trained immunity is an increased abundance of the lipid rafts. The increase in the lipid rafts’ abundance is likely controlled by exNef-mediated suppression of ABCA1 activity (Mukhamedova *et al*., 2019) and may lead to an increase in IGF1R exposure and signaling, which is a key to the mechanism of training (Bekkering *et al*., 2018; Netea *et al*., 2020). Changes in the abundance and properties of the lipid rafts may contribute to trained immunity induced by other factors that modify cellular cholesterol metabolism, including β-glucan, and need to be taken into consideration in future studies.

The contribution of the observed trained immunity to pathogenesis of HIV disease and HIV-associated co-morbidities remains to be investigated. While evolutionary trained immunity has evolved as a protection mechanism, enhanced responses of monocytes/macrophages of HIV-infected individuals to inflammatory stimuli due to trained immunity may explain sustained low-level inflammation when viral load is reduced to undetectable levels by anti-retroviral treatment (ART) (Tincati et al., 2016; Zicari *et al*., 2019). One possible explanation for beneficial features of trained immunity turning into a pathological factor in HIV infection is that protective mechanism of trained immunity cannot deal with long-term influx of pathogenic factors, as happens in HIV infection, where exNef continues to be released from HIV reservoirs (Raymond *et al*., 2019) and bacterial products keep pouring through damaged mucosal tissue (Heron and Elahi, 2017) for a long time after HIV suppression by ART. Thus, contribution of the exNef-mediated training to HIV co-morbidities depends on the duration of mucosal damage and the induced memory. The half-life of circulating monocytes is relatively short (several days), but a possibility of training by exNef of myeloid progenitor cells, a likely mechanism of effects of exNef *in vivo*, may provide a long-lasting memory. Indeed, BMDM from exNef-treated mice showed increased responses to LPS stimulation after 15 days, although they were never directly exposed to exNef. Under this scenario, even complete cure of HIV infection, either by silencing HIV transcription or eliminating integrated HIV sequences from the cells, would still leave infected individuals hyperresponsive to inflammatory stimuli. This may provide a protective effect if the stimuli are acute, but may also put them at risk for inflammation-associated diseases if chronic stimuli are encountered. How long such memory can persist is an important question for future studies, but recent findings by Netea’s group suggested that even a transgenerational transmission of trained immunity is possible (Katzmarski et al., 2021; 2022).

In conclusion, we demonstrated that extracellular vesicles carrying HIV protein Nef may trigger long-lasting changes in inflammatory responses of the host by mechanisms consistent with those of trained immunity. This phenomenon may not only contribute to the persistent chronic inflammation in PLWH, but, considering the demonstrated effects on cholesterol biosynthesis and lipid rafts, may also contribute to comorbidities of HIV infection by other mechanisms. More broadly, trained immunity may also explain the origin of persistent metabolic comorbidities following many viral infections, such as COVID-19.

### Limitations of the study

A limitation of this study is a variability of results between different donors. Although such variability is expected in human samples, much larger number of donors needs to be investigated to obtain quantitative results. Another limitation is that although we establish a contribution of trained immunity to inflammatory responses, the quantitative proportion of this contribution as compared to other mechanisms *in vivo* was not established, neither were different mechanisms compared to each other.

## Acknowledgments

This study was supported by NIH grants R01 HL140977 (M.F. and M.I.B.), R01 HL158305 (D.S. and M.I.B.), P30 AI117970 (M.I.B.). We are grateful to Dr. Christophe Vanpoille for help with characterization of Nef EVs.

## Author contributions

L.D. performed experiments; B.B. performed experiments; N.M.P. analyzed the data; T.P. performed experiments; N.M. performed experiments; D.D. performed experiments; M.F. designed the study; A.H. analyzed the data; A.J.M. interpreted the results; D.S. designed the study, interpreted the results, wrote the manuscript; M.I.B. designed the study, interpreted the results, wrote the manuscript.

## Materials and Methods

### Monocyte preparation, training and differentiation

Buffy coats from healthy donors were obtained from Gulf Coast Blood Center. PBMCs were isolated by density centrifugation using Ficoll-Paque (GE Healthcare) and, after washing 3 times with PBS, cells (9 × 10^6^/ml) were plated in Primaria T-75 flasks or Primaria plates in Dutch modified RPMI (Gibco) supplemented with 1% Pen/Strep (Corning), 10 mM L-Glutamine (Corning) and 10 mM sodium pyruvate (Corning) and incubated in a humidified 37^°^C incubator with 5% CO_2_ for 2 hours to adhere to plastic. Non-adhered cells were removed by washing 3-times with warm PBS (Gibco, DPBS with Mg^2+^ and Ca^2+^), and adhered cells were exposed to exNef or exCont (1 μg/ml protein) for 48 h in the absence or presence of either 20 μM fluvastatin (Sigma, SML 0038) or 10 nM Picropodophyllin (PPP) in Dutch modified RPMI supplemented with 10% Human Serum (HS, CeLLect, #2830349), 1% Pen/Strep, 10 mM L-Glutamine and 10 mM pyruvate. To remove lipid rafts, cells were treated with or 1 mM methyl β-cyclodextrin for 30 min. Cells were washed with warm PBS (with Mg^2+^ and Ca^2+^) and exposed to a fresh complete medium supplemented with M-CSF (20 ng/ml) for the next 6 days. At the end of differentiation, some cells were stimulated with 2 ng/ml lipopolysaccharide (LPS, InvivoGen) for 24 h.

### Flow Cytometry

Phenotypic analysis of monocyte-derived macrophages (MDM) was performed using flow cytometric direct immunostaining. Cells were detached from the plastic with 10 mM EDTA in PBS by scraping. MDM were incubated with Live/Dead Fixable Aqua Dead Cell Stain Kit (Invitrogen) for 20 min on ice, washed with Blocking Buffer (BB, PBS/5% HS/0.0055% NaN_3_) and incubated in BB for 15 min on ice to exclude non-specific binding of Abs. After washing in BB, cells were stained in 100 μl of Brilliant Stain Buffer (BD Pharmingen) (a mixture of mouse anti-human FITC-CD16, Pe-Cy5–CD11b/Mac-1, APC-H7–HLA-DR, BV421– CD38, BV480–CD14, BV786–CD80, APC–TLR4, BV650–CD40, Pe–CD64, Cytofix/Cytoperm Kit) for 30 min on ice. Then cells were washed and incubated in 250 μl of Fixation/Permeabilization Solution for 20 min on ice, washed 2 times in 1 x BD Perm/Wash Buffer, resuspended in 50 μl of BD Perm/Wash Buffer with PerCP/Cy5.5–CD68 (BD Pharmingen), and incubated for 30 min on ice. MDM were washed and resuspended in 50 μl of Staining Buffer (PBS/1% HS/0.0055% NaN_3_) prior to analysis. Flow cytometry was performed on Cytek Aurora (spectral flow cytometry) using SpectroFlo software. SpectroFlo QC beads were used to optimize the cytometer. Unstained, single fluorochrome stained cells and BD CompBeads Compensation Particles (BD Bioscience) were used as Reference Controls to ensure accurate spectral unmixing of the data. Compensation was done in SpectroFlo software (Cytek Biosciences). The results were analyzed with FlowJo software version 10.7.1. The FMO (fluorescence minus one) controls were used for gating on positive cells. To analyze, visualize, and interpret data, equal amounts of MDM treated with exCont or exNef were subjected to tSNE (t-stochastic neighbor embedding) analysis using iteration 1000, perplexity 45, gradient algorithm-Barnes-Hut. Unbiased number of clusters (populations) were determined by Phenograph technique and applied to FlowSom analysis to visualize cytometry data. Phenotypic outcome was considered in the populations where exNef-treated cells prevailed over exCont-treated for 10% or more.

### RNA-seq

Differentiated MDM were washed in PBS and exposed to 2 ng/ml of LPS for 3 h, washed again and used for RNA extraction. Total RNA was isolated using TRIzol (Invitrogen) and cleaned using RNeasy Plus Mini kit (Qiagen). The amount of isolated RNA was measured using Qubit RNA HS Assay Kit (Invitrogen) and Qubit 4 Fluorometer. Sequencing was performed on Illumina HiSeq platform (2 × 150 bp configuration, single index, 50 million reads per sample) by GENEWIZ.

The raw RNA-sequencing reads were QCed using FastQC, and aligned to the latest version of the human genome reference (GRCh38, Dec 2013) using STAR v.2.7.3.c in 2-pass mode with transcript annotations from the assembly GRCh38.79 (Dobin et al., 2013). The alignments were deduplicated, and then sorted using Samtools (Li et al., 2009). Gene counts were estimated using the featureCounts utility of SubRead (Liao et al., 2013). The normalized gene counts were used for differential expression analysis by DeSeq2 (Love et al., 2014). Gene Set Enrichment Analysis (GSEA) toolkit was used to identify pathways enriched in deregulated genes, and to perform leading edge analysis.

### ddPCR

Total RNA isolated for RNA-seq was also subjected to reverse transcription-droplet digital polymerase chain reaction (RT-ddPCR) analysis. Total cDNA from each RNA sample was generated using iScript Reverse Transcription Supermix for RT-qPCR (Bio-Rad). The amount of cDNA used in ddPCR was determined by a pilot experiment using β-actin primers and defined as the amount that will generate Ct value for β-actin approximately equal to 16. Real Time PCR was performed in duplicate using IQ SYBR Green Supermix (Bio-Rad) and CFX96 Real-Time System, Bio-Rad. Gene-specific PCR primer pairs were obtained from OriGene Technologies (Rockville, MD) and used according to company’s protocol. Droplet digital PCR was performed in quadruplicates with QX200 ddPCR EvaGreen Supermix and QXDx AutoDG Droplet Digital PCR System (Bio-Rad). Copy numbers of GAPDH transcripts were used for normalization.

### ATAC-seq

ATAC-seq was performed using ATAC-seq Kit (Cell Biologics, cat# CB6936) according to the instruction manual. In brief: 50,000 cells were spun, washed in cold PBS and resuspended in cold hypotonic buffer (10 mM Tris-HCl, pH 7.4, 10 mM NaCl, 3 mM MgCl_2_ and 0.1% IGEPAL CA-630), centrifuged at 500 g for 10 min at +4^°^C. Nuclei were suspended in transposition reaction mix (25 μl 2 x TD buffer, 2.5 μl Transposome and 22.5 μl nuclease free H_2_0), incubated at 37^°^C for 60 min and purified using Qiagen MinElute kit (Quigen, cat# 28004). Library was generated according to the instructions using the following PCR conditions: 98^°^C for 30 sec.; thermocycling 10 times: 98^°^C for 10 sec., 63^°^C for 30 sec., 72^°^C for 1 min.; hold at 4^°^C. Library was purified using double-sided bead purification method and AMpure XP beads at RT. Library quality and quantity were assessed by qPCR method using NEB Next Library Quant Kit for Illumina (NEB, # E7630) and Qubit. Paired-end sequencing was performed by GENEWIZ on Illumina HiSeq platform (2 × 150 bp configuration, single index, with 50 million reads per sample).

All ATAC-seq sequencing datasets were quality assessed using FastQC, and aligned against the last version of the human reference genome (GRCh38) with bowtie2 (Langmead and Salzberg, 2012) using ‘—very-sensitive’ as parameters. The alignments were subsequently sorted using samtools (Li et al., 2009). The sorted alignments were run though callpeak function of MACS2 (Feng et al., 2011) to produce peak files. Differential analyses between exNef-treated cells and controls were performed using Hurdle model (Cragg, 1971), and a threshold of 10% False Discovery Rate (FDR<0.1) were considered significant.

### Animal experiments

All animal experiments were approved by the Alfred Medical Research Education Precinct (AMREP) animal ethics committee (P1761) and conducted in accordance with the Australian code of practice for the care and use of animals for scientific purposes, as stipulated by the National Health and Medical Research Council of Australia. All mice were housed in a normal light and dark cycle and had *ad libitum* access to food and water. Mice were randomly assigned to treatment and end-point analysis was blinded.

Two groups (6 male mice per group) of C57BL/6J mice (Jackson Laboratories) were administered either exNef (2 μg total protein, I.V.) or control (exGFP) exosomes (2 μg total protein, I.V.), 3 times a week, for a period of 2 weeks as described previously (Mukhamedova *et al*., 2019). At the end of the experiment mice were euthanized, bone marrow cells were washed out from fibia and tibia, plated on Petri dishes, non-adherent cells were washed away while adherent cells (BMDM) were grown for 7 or 15 days in DMEM medium supplemented with 10% FBS and 10% of L929 conditioned medium. Cells were stimulated with LPS (4 h, 200 ng/ml) and medium was collected for analysis of the amount of secreted TNFα by ELISA (Invitrogen).

### Cholesterol biosynthesis

Cells were plated in 24-well plates and incubated with exNef or exCont for 48 h, washed and incubated with [^3^H] acetate (final radioactivity of 3.7 MBq/ml) in serum-free medium supplemented with 0.1% BSA for 2 hrs. Cells were washed twice with PBS, collected in water, lipids were extracted and cholesterol isolated by TLC as described previously (Fu et al., 2004) and counted in β-counter (Hidex).

### Plasma membrane protein isolation

Plasma membrane proteins were isolated using the Minute™ kit from Invent Biotechnologies (Plymouth, MN) following manufacturer’s protocol. For lipid raft isolation, membrane fraction was isolated and subjected to centrifugation in Iodixanol density gradient as described previously (Mukhamedova *et al*., 2019; Mukhamedova et al., 2020).

### Western blotting

Samples were analyzed by automated Western immunoblotting using the JessTM Simple Western system (BioTechne, San Jose, CA). For analysis of Flotillin 1 and β actin, a 20-230 kDa Jess separation module SM-W004 was used. Plasma membrane (60 mg per sample) was homogenized in the Triton X-100 RIPA lysis buffer (250 μL, ThermoFisher) and cleared by centrifugation at 5,000 x g. Three μL of lysate (1 μg/μL protein) was mixed with 1 μL of Fluorescent 5X Master mix (BioTechne) in the presence of fluorescent molecular weight markers and 400 mM dithiothreitol (BioTechne). This preparation was denatured at 95°C for 5 min. Molecular weight ladder and proteins were separated in capillaries through a separation matrix at 375 volts. A BioTechne proprietary photoactivated capture chemistry was used to immobilize separated viral proteins on the capillaries. Capillaries with immobilized proteins were blocked with KPL Detection Block (5X) (SeraCare Life Sciences, Gaithersburg, MD, cat. #5920-0004) for 60 min, and then incubated with primary anti-hFlotillin 1 goat polyclonal antibody (BioTechne) or anti-hβ-actin mouse monoclonal antibody (R&D) for 60 min. After a wash step, HRP-conjugated anti-goat (cat. #043-522-2) or near infrared (NIR) fluorescent dye-conjugated anti-mouse (cat. #043-821) secondary antibody from BioTechne was added for 30 min to capillaries. The chemiluminescent revelation was established with peroxide/luminol-S (BioTechne). NIR imaging was obtained using Jess fluorescent detection channel. Digital image of chemiluminescence or fluorescence of the capillary was captured with Compass Simple Western software (version 5.1.0, Protein Simple) that calculated automatically heights (chemiluminescence or fluorescence intensity), area, and signal/noise ratio. Results were visualized as electropherograms representing peak of chemiluminescence or fluorescence intensity and as lane view from signal of chemiluminescence or fluorescence detected in the capillary. A total protein assay using Total Protein detection module DM-TP01 and Replex Module RP-001 was included in each run to quantitate loading. Samples were analyzed at least 2 times to ensure consistency of the results.

## Supplemental information

**Figure S1. RNA-seq analysis**. Monocytes from 8 donors were treated with exNef or exCont during the first 48 h of differentiation, washed and cultured for additional 6 days. Cells were treated with LPS, RNA was isolated and analyzed by RNAseq. **A** - HeatMap of the most deregulated genes. **B** - Gene set enrichment analysis showing enrichment in inflammatory and cytokine pathways. **C** - Leading edge analysis shows that a number of cytokine genes, including IL-2, IL-3, IL-6, IL-11, CXCL8 and TNFα, drive the changes of several inflammatory pathways.

**Figure S2. Analysis of lipid rafts**. Distribution of [^3^H]cholesterol (**A**) and flotillin-1 (**B**) after fractionation of plasma membranes by ultracentrifugation in a density gradient. BMDM derived from BM of animals treated with either exNef or exGFP (7 days after isolation) were labelled *in vitro* with [^3^H]cholesterol. Membrane fraction was isolated and subjected to centrifugation in Iodixanol density gradient as described previously (Mukhamedova *et al*., 2019; Mukhamedova *et al*., 2020). [^3^H]cholesterol content was determined by β-counting, and flotillin-1 content was determined by densitometry of flotillin-1 bands after subjecting each fraction to PAGE followed by Western blotting. Fractions were collected from the top of the gradient (1st fraction is the least dense fraction). AUC – area under the curve. ***p<0.001 (difference between the curves, paired t-test).

**Figure S3. t-SNE analysis of MDM**. Equal amounts of MDM from two donors (panels A, B – one donor, and C, D – another donor) exposed to exCont or exNef were subjected to t-SNE (t-stochastic neighbor embedding) analysis of multiparametric flow staining for inflammatory markers (see Fig. 2 for details). Unbiased number of clusters (populations) were determined by Phenograph technique and applied to FlowSom analysis to visualize cytometry data. **A, C** - Results are shown for ungated population, and after gating on exCont or exNef exposed cells. **B, D** - Phenotypic outcome was evaluated in the populations where exNef-treated cells prevailed over exCont-treated by 50% or more. Top left panel shows distribution of these cell populations, and the table underneath presents the frequency of parent population for each cluster. The diagrams on the right show expression of indicated markers on the cells of each color-coded population.

**Table S1.** **Genes that changed localization between open and closed chromatin regions (comparison of results of ATACseq with cells from 4 donors treated with exNef or exCont).**

| **Region ID** | **Gene** | **P-value** | **FDR** | **Fold change (exNef vs exCont)<sup>1</sup>** | **Mean exNef** | **Mean exCont** | **Pathway** |
| --- | --- | --- | --- | --- | --- | --- | --- |
| chr2:169012492-169012682 | ABCB11 | 0.00 | 0.00 | 23.45 | 2.54 | 0.11 | Cholesterol Metabolism |
| chr22:17107426-17107604 | IL17RA | 0.00 | 0.00 | 21.74 | 2.36 | 0.11 | Inflammatory Response |
| chr4:109702595-109702794 | CASP6 | 0.00 | 0.00 | 19.84 | 2.15 | 0.11 | Inflammatory Response |
| chr11:63681662-63681740 | RTN3 | 0.00 | 0.04 | 7.84 | 1.55 | 0.20 | Inflammatory Response |
| chr18:47930279-47930434 | SMAD2 | 0.00 | 0.07 | 2.34 | 6.32 | 2.71 | Cholesterol Metabolism |
| chr7:21554070-21554257 | DNAH11 | 0.00 | 0.03 | -2.15 | 3.05 | 6.58 | Inflammatory Response |
| chr11:121292824-121292965 | SC5D | 0.00 | 0.00 | -17.84 | 0.12 | 2.13 | Cholesterol Metabolism |

<sup>1</sup>Positive values indicate increased presentation in open chromatin after exNef treatment, whereas negative values - decreased presentation.

