## Supplemental Figure 1 for "Extracellular vesicles carrying HIV-1 Nef induce trained immunity in myeloid cells"

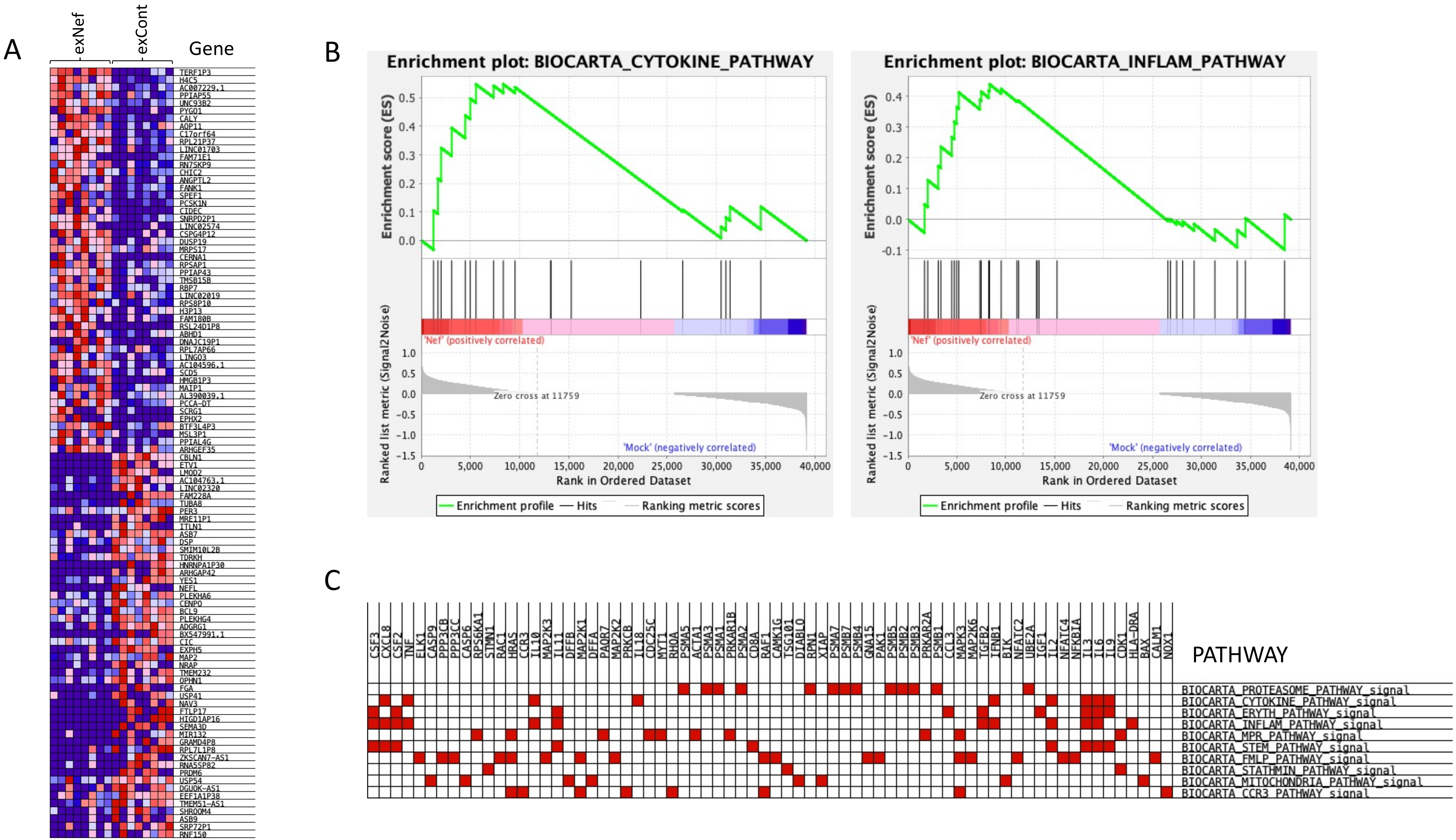


**Figure S1. RNAseq analysis.** Monocytes from 8 donors were treated with exNef or exCont during the first 48 h of differentiation, washed and cultured for additional 6 days. Cells were treated with LPS, RNA was isolated and analyzed by RNAseq. **A** - HeatMap of the most deregulated genes. **B** - Gene set enrichment analysis showing enrichment in inflammatory and cytokine pathways. **C** - Leading edge analysis shows that a number of cytokine genes, including IL-2, IL-3, IL-6, IL-11, CXCL8 and TNFα, drive the changes of several inflammation-related pathways.
