## Supplemental Figure 2 for "Extracellular vesicles carrying HIV-1 Nef induce trained immunity in myeloid cells"

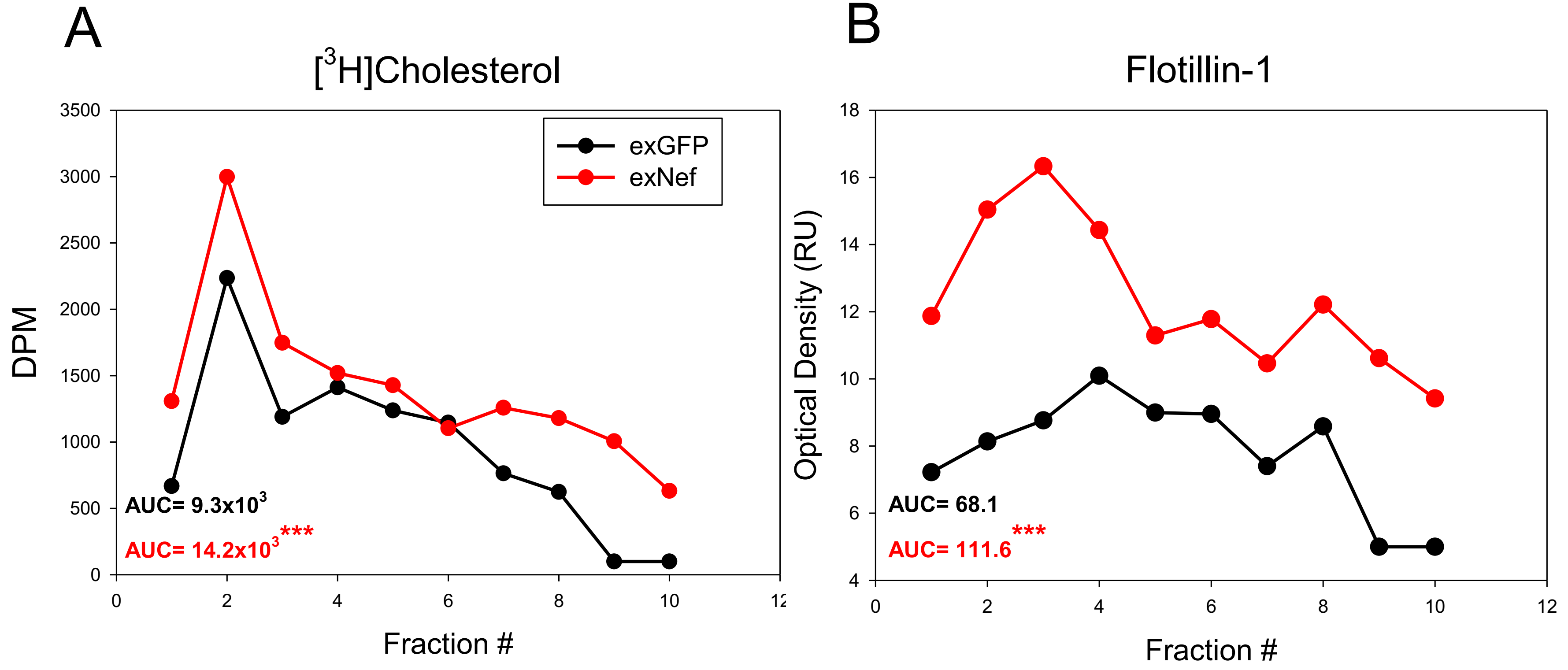


**Figure S2. Analysis of lipid rafts.** Distribution of $[^3H]$cholesterol (**A**) and flotillin-1 (**B**) after fractionation of plasma membranes by ultracentrifugation in a density gradient. BMDM derived from BM of animals treated with either exNef or exGFP (7 days after isolation) were labelled *in vitro* with $[^3H]$cholesterol. Membrane fraction was isolated and subjected to centrifugation in Iodixanol density gradient as described previously (Mukhamedova *et al.*, 2019; Mukhamedova *et al.*, 2020). $[^3H]$cholesterol content was determined by β-counting, and flotillin-1 content was determined by densitometry of flotillin-1 bands after subjecting each fraction to PAGE followed by Western blotting. Fractions were collected from the top of the gradient (1st fraction is the least dense fraction). AUC – area under the curve. ***$p<0.001$ (difference between the curves, paired t-test).
