## Supplementary figures and images for "Extracellular vesicles carrying HIV-1 Nef induce trained immunity in myeloid cells"

### Supplemental Figure 3

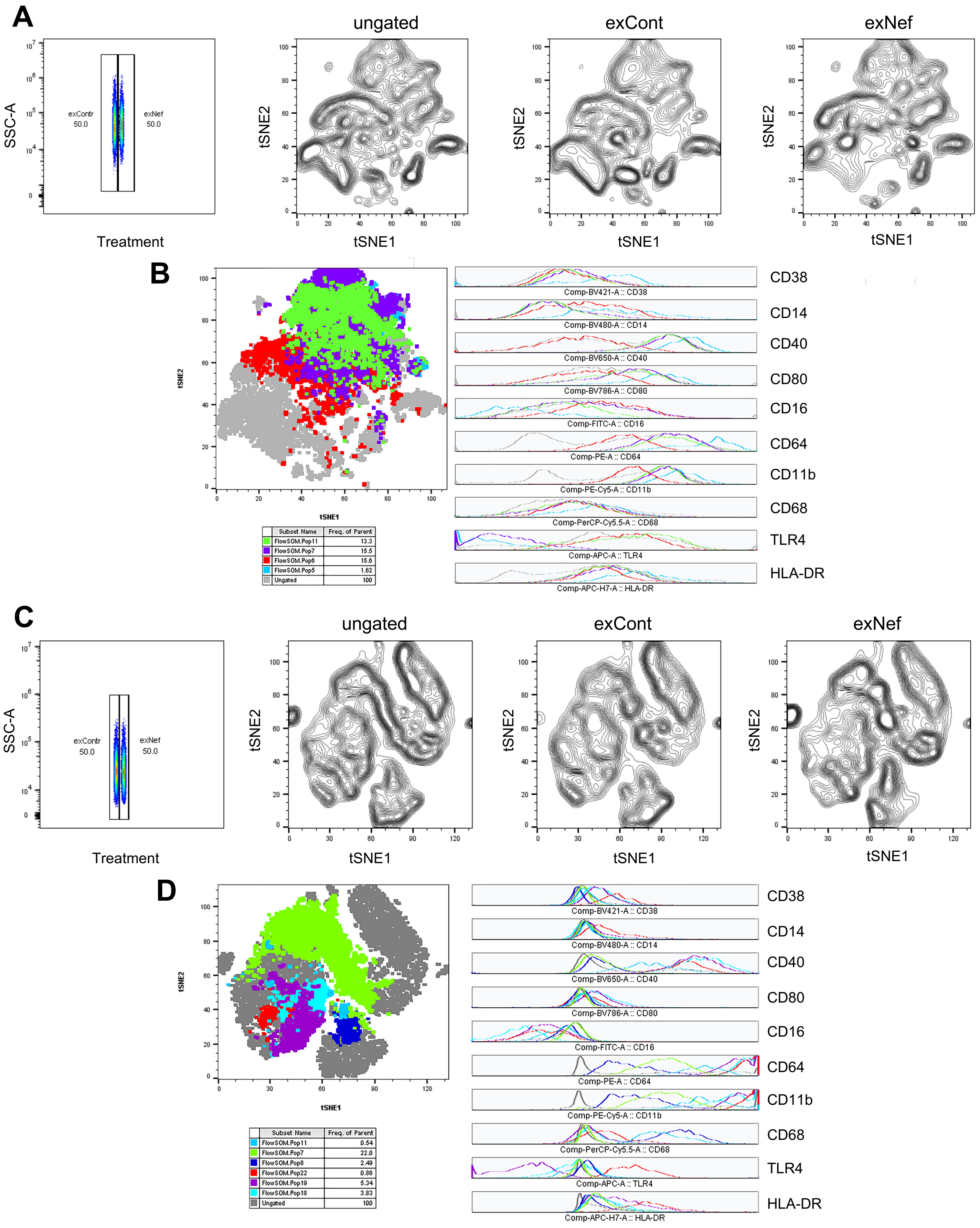


**Figure S2. t-SNE analysis of MDM from 2 additional donors.**
